# NeCS-Flow: Necrosis-Guided Contrast-Enhanced MRI Synthesis with Rectified-Flow Refinement

**DOI:** 10.64898/2026.10.21.760821

**Authors:** Eunate Alzaga Goñi, Walter Zhao, Rhea Adams, Shahrzad Moinian, Yanfei Xiong, Tianhe Wu, Kewei Yan, Katie Shpanskaya, Anil Vasireddi, Jessica Houk, Evan Calabrese, Dan Ma

## Abstract

**Background:** Gadolinium-based contrast agents raise concerns about tissue deposition, adverse reactions, and increased costs.

**Purpose:** To develop and validate a deep learning pipeline for gadolinium-free post-contrast T1w brain MRI synthesis incorporating necrotic scaling and Posterior-Mean Rectified Flow (PMRF) refinement, and to evaluate diagnostic utility through a blinded multi-reader study.

**Study Type:** Retrospective.

**Population:** 454 patients with adult-type gliomas (grade 2-4) from four publicly available datasets. Training used 374 patients; internal testing used 60 held-out patients; external validation used 20 patients.

**Field Strength/Sequence:** 1.5-3.0 T; pre-contrast T1w, T2w, and FLAIR as inputs; post-contrast T1w as synthesis target.

**Assessment:** A three-stage pipeline comprising nnU-Net Necrotic core segmentation, necrotic-scaling-guided U-Net Contrast Synthesis, and PMRF perceptual refinement (NeCS-Flow) was evaluated using whole-brain and tumor-region similarity metrics, enhancing tumor Dice scores, perceptual quality metrics, and a blinded three-reader study assessing diagnostic accuracy, enhancement detection, image quality, and real-versus-synthetic discriminability.

**Statistical Tests:** Paired t-tests and two one-sided tests for model comparison; Fisher’s exact test and two-sample t-tests for reader study comparisons, with p < 0.05 considered significant.

**Results:** Necrotic scaling consistently improved real versus synthetic enhancing tumor Dice internally (HGG: 0.793 to 0.816) and externally (HGG: 0.465 to 0.612). NeCS-Flow outperformed two benchmark architectures on all quantitative metrics in internal and external testing, although absolute enhancing tumor accuracy declined for all models on the external cohort. PMRF refinement improved perceptual realism across gliomas, reducing both FID and KID. Readers distinguished synthetic from real images only modestly above chance (overall accuracy 61.7%), and no significant differences between real and synthetic images were found for any clinical metric.

**Data Conclusion:** NeCS-Flow synthesizes perceptually realistic and diagnostically equivalent post-contrast T1w brain MRI, with necrotic scaling improving enhancing tumor delineation and PMRF refinement improving perceptual realism across gliomas. Performance on an external cohort was preserved relative to benchmark methods, though reduced in absolute terms.

## 1. INTRODUCTION

Magnetic resonance imaging (MRI) is essential for brain tumor diagnosis, treatment planning, and therapeutic monitoring due to its superior soft tissue contrast compared to alternative modalities like computed tomography. Gadolinium-based contrast agents (GBCAs) highlight tumor boundaries and blood-brain barrier disruption, aiding surgical planning and treatment optimization (1, 2). However, gadolinium deposition in the brain and bone tissue following GBCA administration (3, 4) raises concerns about long-term effects of repeated exposure. Modern macrocyclic GBCAs have improved safety profiles compared with older linear agents, but concerns about gadolinium retention have motivated continued interest in gadolinium-free contrast-enhanced image synthesis methods (5).

Standard clinical brain tumor MRI protocols include multiple pre-contrast sequences that provide complementary information (6), which deep learning models can leverage to synthesize post-contrast images (7–12). However, two important limitations remain unresolved: 1) spatial fidelity of tumor enhancement, and 2) perceptual fidelity. First, synthesized images often struggle to faithfully reproduce the spatial distribution of tumor enhancement. Because tumor voxels are underrepresented during training, synthesis models prioritize the more abundant healthy brain tissue, resulting in less accurate representation of tumor regions. Recent approaches have attempted to address this limitation using local loss functions (8) and semantic feature-extraction modules (12). However, these methods primarily focus on the tumor as a whole and do not explicitly account for the heterogeneous spatial patterns of contrast enhancement within tumors. As a result, enhancement may be underrepresented or spatially mislocalized, particularly in tumors with complex enhancement patterns. Second, accurate voxelwise reconstruction does not necessarily translate into perceptually realistic images. Networks optimized with voxelwise losses regress toward the conditional mean, producing over-smoothed outputs that can score well on distortion metrics yet remain perceptually distinguishable from real images (13, 14). Despite the importance of perceptual realism for clinical adoption, it is seldom assessed through formal reader studies (15).

Our study aims to address both limitations by developing a deep learning pipeline comprising two complementary components. First, a necrotic-scaling stage uses a necrotic core label to spatially weight the input images, incorporating the physiological relationship between necrosis and surrounding blood–brain barrier disruption (16, 17) directly into the synthesis process. Second, a Posterior-Mean Rectified Flow (PMRF) (18, 19) refinement stage maps the over-smoothed posterior-mean prediction from the synthesis network onto a realistic sample from the image distribution, restoring tissue and enhancement texture lost to voxelwise averaging. We refer to this pipeline as NeCS-Flow. We hypothesize that necrotic scaling will maintain image quality comparable to standard synthesis while improving the accuracy of predicted contrast enhancement where present, and that PMRF will improve perceptual realism without compromising quantitative fidelity, as assessed through a blinded multi-reader study. The pipeline is evaluated across two adult brain tumor types, high-grade gliomas (HGGs; WHO grade 4) and low-grade gliomas (LGGs; WHO grades 2-3), with external test data used to determine whether the improvements in enhancing tumor regions conferred by necrotic scaling generalize beyond the training institutions.

## 2. MATERIALS AND METHODS

### 2.1 Study Cohorts and Datasets

Data from four publicly available datasets were used: three for model development and internal evaluation, and one for external evaluation. The training set comprised 374 patients: 300 from the University of Pennsylvania de novo glioblastoma (GBM) collection (UPENN-GBM, 2006–2018) (20), 40 from the UCSF Preoperative Diffuse Glioma MRI dataset (UCSF-PDGM, 2015–2021) (21), and 34 from the Cancer Genome Atlas low-grade glioma collection (BraTS-TCGA-LGG) (22). Dataset characteristics are summarized in Table 1. UPENN-GBM cases were all HGGs (WHO grade 4), and UCSF-PDGM and BraTS-TCGA-LGG cases were all LGGs (WHO grades 2-3). UCSF-PDGM grade 4 cases were excluded to maintain this grouping. Training cases were selected at random from each dataset. LGG numbers were limited by availability.

**Table 1:** Dataset characteristics.

| Parameter | UPENN-GBM | UCSF-PDGM | BraTS-TCGA-LGG | OpenNeuro ds007045 |
| --- | --- | --- | --- | --- |
| Number of patients | 330 | 56 | 48 | 20 |
| Mean age ± standard deviation | 63 ± 12 years | 42 ± 14 years | 44 ± 14 years | 59 ± 11 years |
| Sex | 197 males, 133 females | 31 males, 25 females | 24 males, 24 females | 7 males, 13 females |
| Diagnosis |  |  |  |  |
| Glioblastoma | 330 | 0 | 0 | 20 |
| Astrocytoma | 0 | 45 | 12 | 0 |
| Oligodendroglioma | 0 | 11 | 15 | 0 |
| Mixed glioma / Not specified | 0 | 0 | 21 | 0 |
| IDH Status |  |  |  |  |
| IDH-mutant | 0 | 46 | 1 | 0 |
| IDH-wildtype | 330 | 10 | 0 | 20 |
| Not specified | 0 | 0 | 47 | 0 |

UPENN-GBM includes de novo GBM patients imaged on 3.0-T Siemens and GE scanners. UCSF-PDGM includes adults with histopathologically confirmed WHO grade 2-4 diffuse gliomas imaged pre-surgically at 3.0 T (GE Discovery 750). Both provide T2w, FLAIR, DWI, and pre- and post-contrast T1w; BraTS-TCGA-LGG provides the same sequences except DWI.

The internal test set comprised 60 held-out patients, balanced across tumor type: 30 HGGs from UPENN-GBM and 30 LGGs (16 from UCSF-PDGM and 14 from BraTS-TCGA-LGG). Additional testing was performed on 20 GBM cases from an external, multi-institutional glioblastoma dataset (OpenNeuro ds007045) (23). No publicly available external LGG dataset containing the required sequences was identified. The only exclusion criterion applied across datasets was severe imaging artifacts, resulting in the exclusion of one UPENN-GBM case from the internal test set.

### 2.2 Image Preprocessing and Tumor Labels

Each dataset was used as publicly released, with standardized preprocessing already applied to it. UPENN-GBM, UCSF-PDGM, and BraTS-TCGA-LGG were skull-stripped, co-registered, and resampled to 1 mm³ isotropic resolution, with tumor segmentations generated automatically and refined by expert neuroradiologists. All three datasets followed the BraTS labeling convention, delineating the necrotic core, contrast-enhancing region, and peritumoral FLAIR hyperintensity.

### 2.3 Three-Stage Model Architecture

This section presents an overview of the three-stage architecture with model details in the following sections. NeCS-Flow was implemented as a three-stage cascaded framework (Figure 1). In Stage 1, an nnU-Net model (24) segments the tumor necrotic core from pre-contrast sequences, identifying regions where enhancement is not physiologically expected and providing the core used for scaling at inference. This segmentation constructs a spatially adaptive scaling map, applied multiplicatively to pre-contrast T1w, T2w, and FLAIR signal intensities within the necrotic region before synthesis, with weights increasing smoothly from 1.0 at the necrotic boundary to 2.0 at its core (see Supplementary Material, Section 2), designed to reduce underprediction and mislocalization of enhancement. In Stage 2, the scaled inputs are passed to a 3D fully convolutional U-Net (7, 25) (Python 3.10, TensorFlow 2.10) that synthesizes the post-contrast T1w image, and in Stage 3 this output is refined by a PMRF (19) model to restore realistic texture and mitigate over-smoothing. Two variants were compared throughout: a baseline without necrotic scaling, and the full NeCS-Flow model.

**Figure 1.**
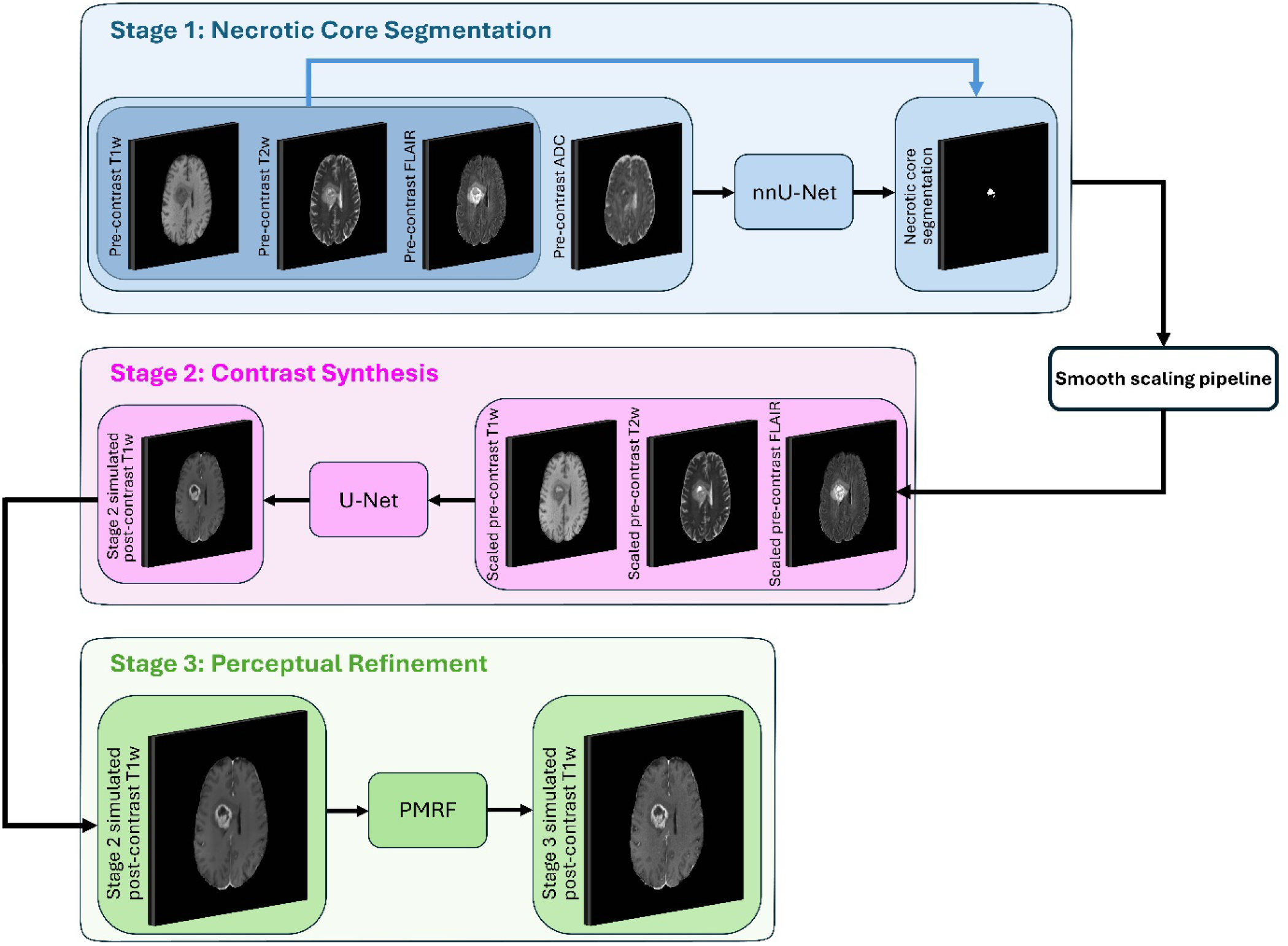
Overview of the NeCS-Flow three-stage contrast synthesis pipeline. In Stage 1, an nnU-Net model segments the necrotic core from pre-contrast T1w, T2w, FLAIR images and ADC/AD maps. The resulting segmentation drives a smooth scaling pipeline that up-weights necrotic core voxels in the pre-contrast inputs. In Stage 2, the scaled pre-contrast T1w, T2w, and FLAIR images are passed to a 3D fully convolutional U-Net to synthesize the post-contrast T1w image. In Stage 3, the U-Net output is refined by a PMRF model to restore realistic image texture and mitigate over-smoothing.

### 2.4 Model Training and Ablation Experiments

The Stage 1 nnU-Net was trained on pre-contrast T1w, T2w, FLAIR images, and apparent diffusion coefficient (ADC)/axial diffusivity (AD) maps. T2w and FLAIR highlight regions of high water content including infiltrative tumor, necrosis and peritumoral edema (26, 27), while elevated ADC reflects the increased water diffusivity characteristic of necrotic tissue (28) and decreased ADC reflects highly cellular regions of active tumor or cytotoxic edema (29, 30). The model was trained in the 3D full-resolution configuration with five-fold cross-validation using nnU-Net’s self-configuring preprocessing and network design.

The synthesis U-Net was trained on pre-contrast T1w, T2w, and FLAIR to synthesize post-contrast T1w images, using 3D patches (100×100×100 voxels) on two NVIDIA Tesla V100 GPUs (32 GB each), with a 90/10 training/validation split. Prediction uncertainty was quantified via Monte Carlo dropout at test time (50 forward passes per case), with the voxelwise standard deviation across samples used as the uncertainty map.

The PMRF refinement stage was trained for 50 epochs to map the U-Net posterior-mean prediction to the ground-truth post-contrast image via rectified flow (18, 19). Pre-computed U-Net predictions served as the flow source, perturbed with Gaussian noise (σ = 0.1). A 3D time-conditioned U-Net was trained with a flow-matching objective using the Adam optimizer (learning rate 1×10⁻⁴), batch size 4, Z-score intensity normalization, and a 90/10 training/validation split (see Supplementary Material, Section 3.3). To assess the contribution of individual input sequences, an ablation study was performed across four input configurations: (1) T1w + T2w + FLAIR, (2) T1w + T2w, (3) T1w + FLAIR, and (4) T2w + FLAIR.

For comparison, two state-of-the-art benchmark architectures, HRNet (8) and DDResUNet (31), were also implemented, trained on identical splits and input sequences and evaluated with identical metrics, including on the external HGG test set, for direct comparison of generalization behavior (Figure 2, Table 3).

**Figure 2.**
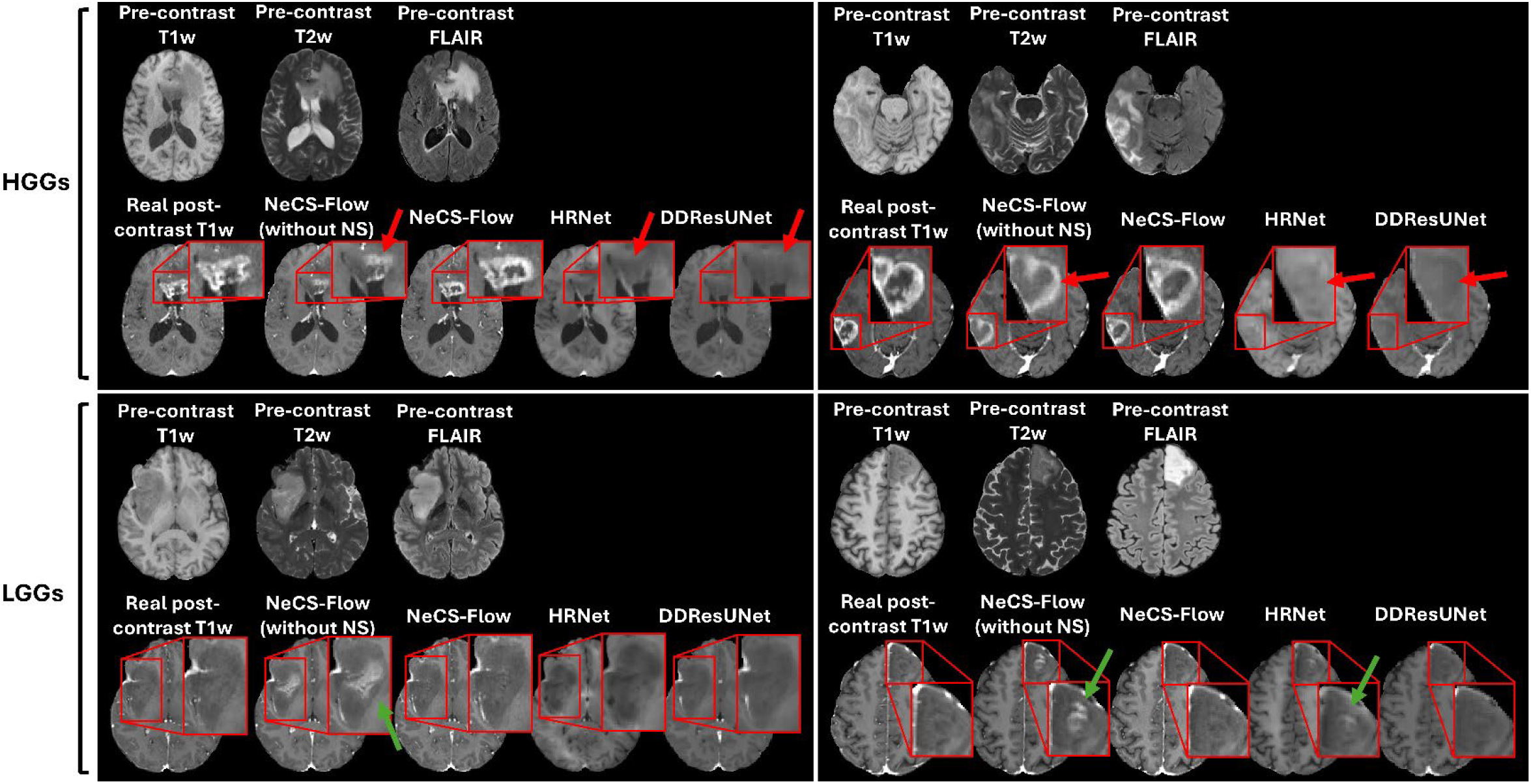
Qualitative comparison of predicted post-contrast T1w images across methods for representative HGG and LGG cases. For each case, the pre-contrast T1w, T2w, and FLAIR inputs are shown in the top row, while the bottom row shows the real post-contrast T1w image, the predicted post-contrast T1w image without necrotic scaling, the proposed method with necrotic scaling (NeCS-Flow), and two state-of-the-art baseline methods (HRNet and DDResUNet). Red arrows indicate incomplete or blurred enhancing rims in the HGG cases; green arrows indicate subtle false-positive enhancement in the LGG cases.

**Table 2:**
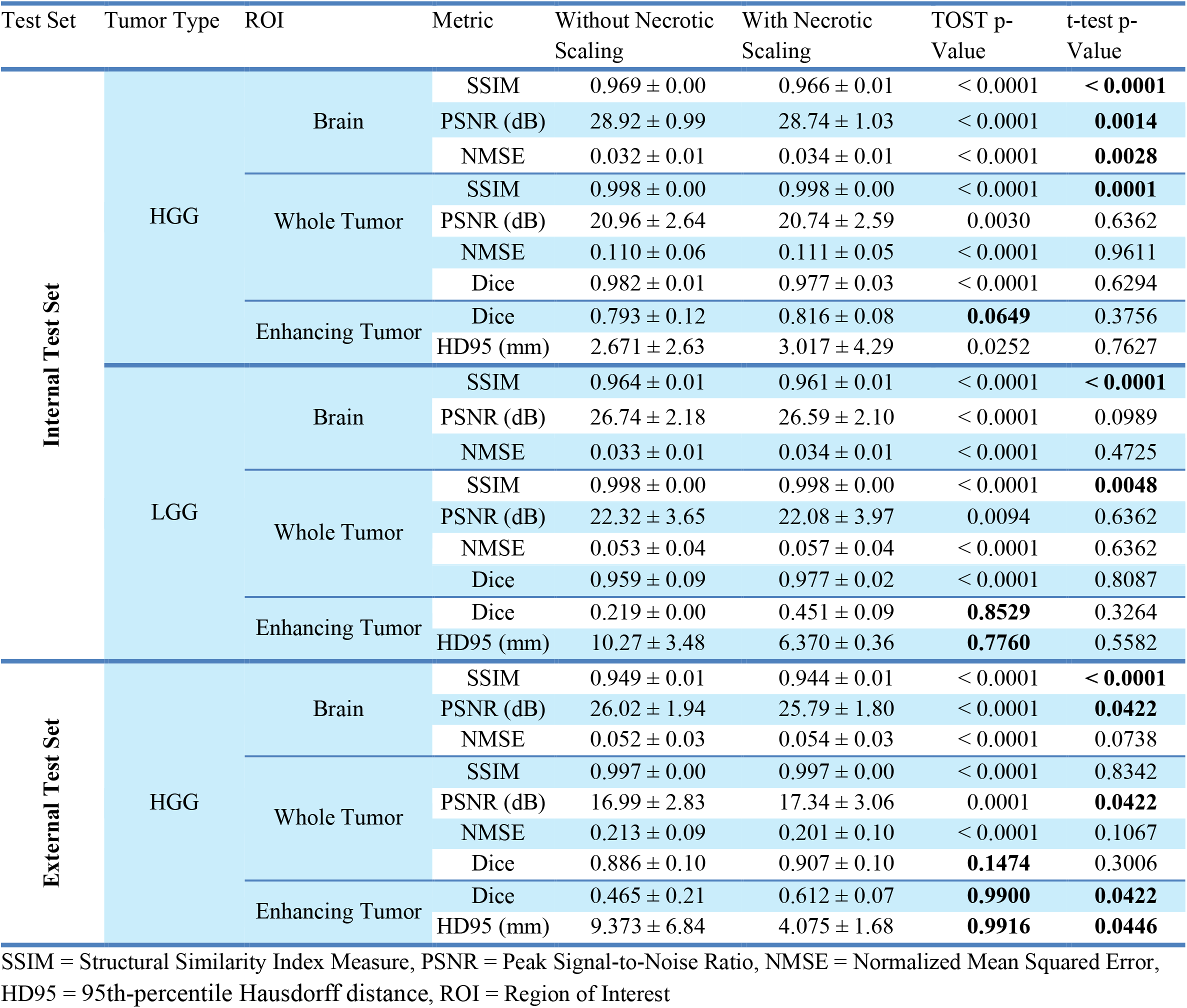
Quantitative results of test data stratified by tumor type.

**Table 3:**
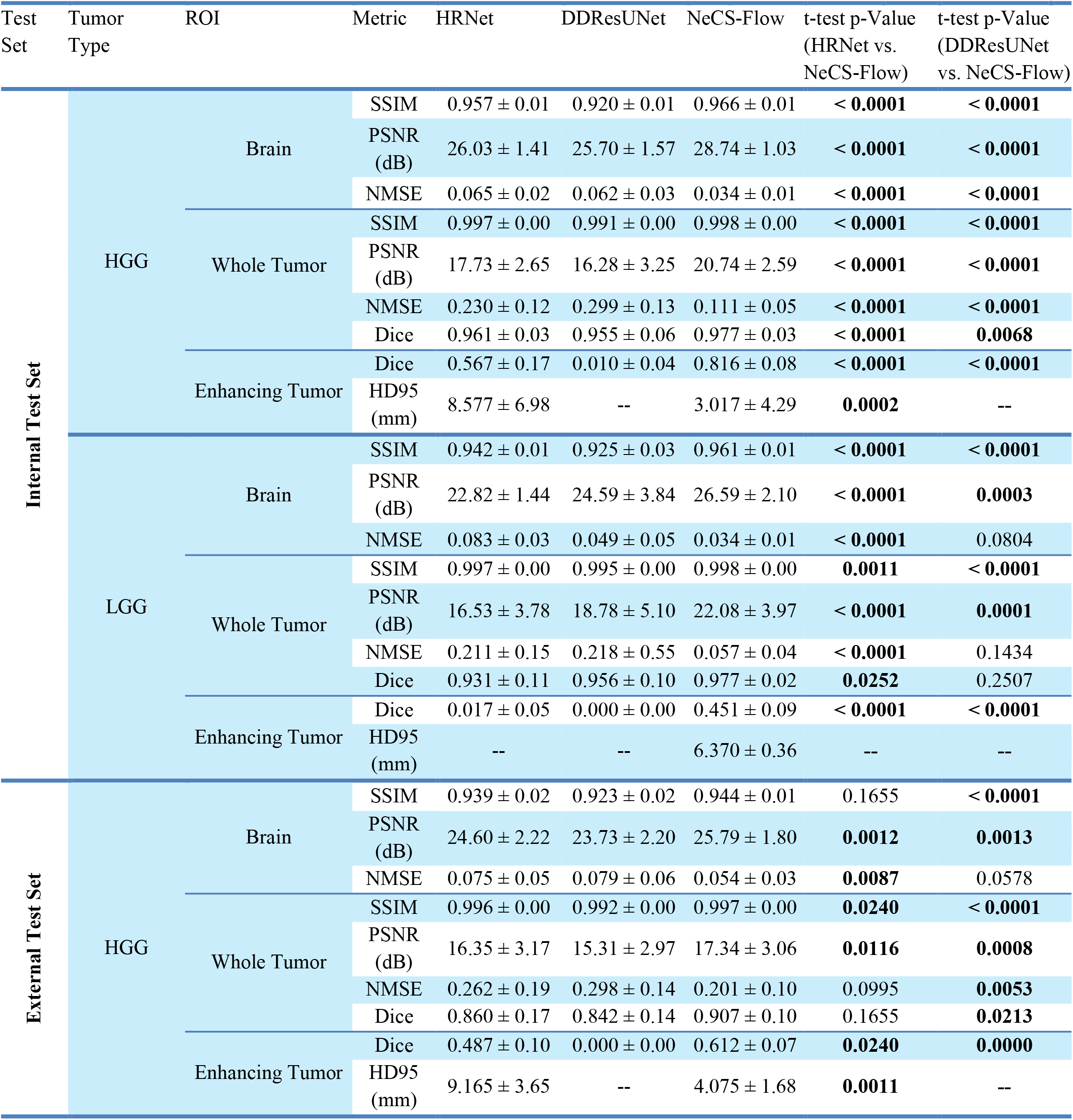
Quantitative results of test data compared with SOTA models.

### 2.5 Blinded Multi-Reader Study

A blinded, multi-reader study was conducted to evaluate the clinical utility and realism of the synthetic contrast-enhanced images. Three readers participated: two board-certified neuroradiologists with 3 and 2 years of attending experience (R1 and R2, respectively), and a neuroradiology fellow (R3). Readers independently reviewed images on a web-based platform (PACSbin), recording responses in REDCap. For each of the 60 internal test cases, the real and synthetic post-contrast images were assessed in parallel by two different readers, one reviewing the real image and the other the synthetic image. Each reader assessed 40 images, with an equal distribution across tumor types, presented in random order.

Assessment spanned four domains. For diagnostic utility, readers classified tumor type and rated diagnostic confidence (1 = not confident at all, 5 = extremely confident). For enhancement quality, they indicated the presence of pathological enhancement (yes/no) and rated enhancement realism (1 = clearly artificial, 5 = very realistic). For image quality, they identified interpretation-affecting artifacts (yes/no) and rated overall image quality (1 = very poor, 5 = excellent). Finally, readers judged whether each case was real or synthetic, enabling assessment of whether synthetic images were perceptually distinguishable from real ones (see Supplementary Material, Section 4, for the full protocol).

### 2.6 Quantitative and Statistical Analysis

Models were evaluated on the 60-patient internal test set and the 20-patient external dataset, with means and standard deviations computed for all metrics. Similarity metrics (SSIM, PSNR) and an error metric (NMSE) were computed between synthetic and real post-contrast T1w images for both the whole-brain and whole-tumor regions. Dice scores quantified spatial overlap for whole-tumor and enhancing-tumor regions, and enhancing-tumor boundary accuracy was assessed using the 95th-percentile Hausdorff distance (HD95). Enhancing and whole-tumor regions in real and synthetic images were segmented using a separate nnU-Net model, distinct from the Stage 1 necrotic-core segmentation model. Models were compared using two complementary statistical tests. Paired t-tests assessed whether necrotic scaling produced a significant difference in each metric, while two one-sided tests (TOST) assessed equivalence for whole-brain and tumor image quality metrics, with equivalence margins of δ = 0.01 (SSIM), 1.0 dB (PSNR), 0.05 (NMSE), 0.05 (Dice), and 2.00 mm (HD95), reflecting thresholds below which differences are clinically negligible. The proposed method was additionally compared with each benchmark architecture on the internal and external test sets using paired t-tests. P-values across metrics and tumor-type strata were adjusted using the Benjamini-Hochberg procedure. To assess the perceptual realism of Stage 2 versus Stage 3 predictions, Fréchet Inception Distance (FID) and Kernel Inception Distance (KID) were computed at the population level for each tumor type. For the reader study, categorical metrics (diagnostic accuracy, enhancement detection accuracy, artifact detection) were compared using Fisher’s exact test, and continuous metrics (diagnostic confidence, enhancement realism, image quality) using two-sample t-tests.

## 3. RESULTS

### 3.1 Qualitative Evaluation

Figure 2 illustrates the effect of necrotic scaling on synthesis quality across representative cases from both tumor types. In the HGG cases, the model without necrotic scaling produces incomplete or blurred enhancing rims (red arrows), whereas necrotic scaling yields sharper, more complete ring enhancement that closely matches the real post-contrast image. In the LGG cases, the model without necrotic scaling tends to produce subtle false-positive enhancement (green arrows), which is suppressed with necrotic scaling. Compared with the benchmark models, NeCS-Flow better preserves the extent and continuity of enhancement in the HGG examples, which the benchmarks underestimate.

Figure 3 illustrates the qualitative effect of the PMRF refinement stage (Stage 3) relative to the U-Net synthesis output (Stage 2) across both tumor types, alongside voxelwise prediction uncertainty maps and population-level perceptual quality metrics. The Stage 3 output consistently exhibits sharper tissue boundaries, more natural gray-white matter contrast, and finer sulcal detail than the Stage 2 predictions, reflected quantitatively in reductions in both FID and KID: for HGGs, FID decreased from 66.75 to 40.06 and KID from 0.0310 to 0.0083; for LGGs, FID decreased from 80.65 to 53.00 and KID from 0.0435 to 0.0159. The uncertainty maps show the highest prediction variance in and around the enhancing tumor. The HGG cases also demonstrate accurate reproduction of diagnostically relevant patterns, including two satellite enhancing lesions (yellow arrows, first case) and leptomeningeal enhancement (second case). Both LGG cases are non-enhancing, illustrating correct suppression of enhancement in the absence of contrast uptake.

**Figure 3.**
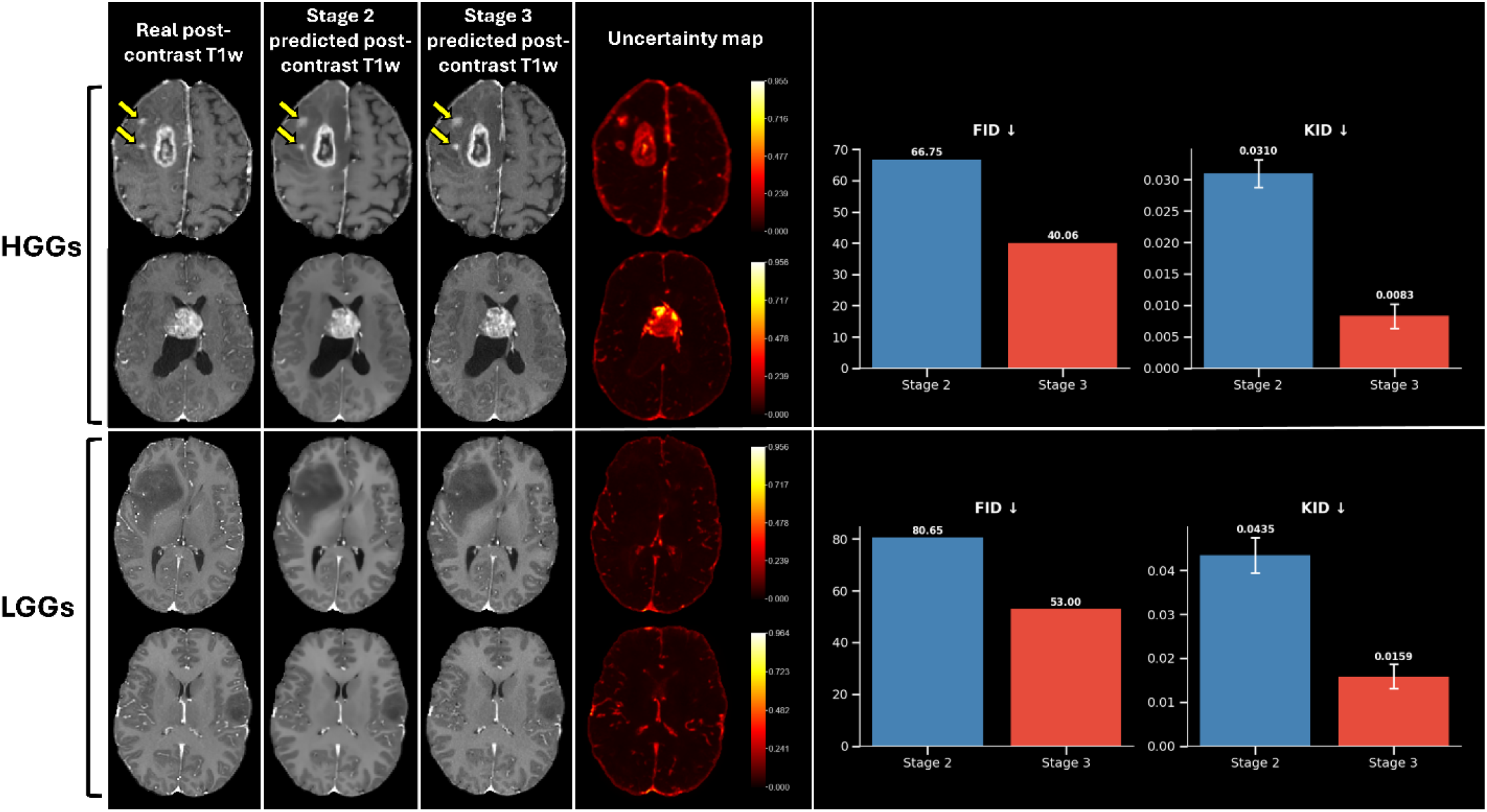
Qualitative comparison of Stage 2 (U-Net) and Stage 3 (PMRF) predicted post-contrast T1w images alongside prediction uncertainty maps and population-level perceptual quality metrics for HGG and LGG cases. For each tumor type, two representative subjects are shown alongside the corresponding real post-contrast T1w image. The uncertainty map represents the voxelwise standard deviation across 50 Monte Carlo dropout samples from the Stage 2 U-Net. Bar charts show FID and KID for Stage 2 and Stage 3 predictions. Yellow arrows in the first HGG case indicate two satellite enhancing lesions accurately reproduced by both Stage 2 and Stage 3 outputs.

### 3.2 Quantitative Evaluation

The input sequence ablation study for the Stage 3 output, conducted on the internal test set, is summarized in Figure 4. Across the four configurations evaluated, T2w+FLAIR showed the largest degradation in whole-brain metrics and the lowest whole-tumor PSNR and NMSE. T1w+T2w performed closest to the full three-sequence reference for whole-brain metrics, while T1w+FLAIR performed closest within the tumor region.

**Figure 4.**
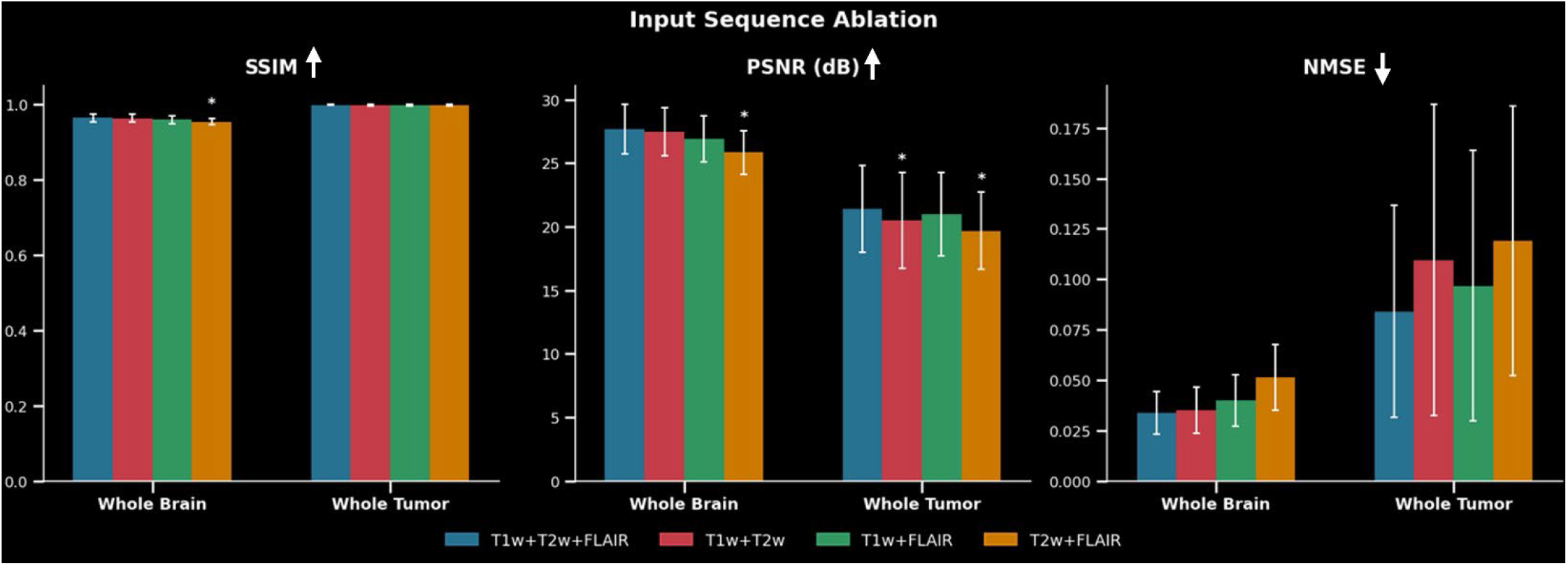
Input sequence ablation study comparing synthesis performance across four input configurations: T1w+T2w+FLAIR, T1w+T2w, T1w+FLAIR, and T2w+FLAIR. Bar charts show mean and standard deviation of SSIM, PSNR, and NMSE for whole-brain and whole-tumor regions. Asterisks indicate configurations that fell outside the predefined equivalence margins relative to the full three-sequence reference model.

Necrotic core segmentation performance of the Stage 1 nnU-Net model is illustrated in Figure 5. For HGGs, the model achieved an average Dice of 0.770 ± 0.197, showing good overlap between ground truth and predicted necrotic regions. For LGGs, the average Dice was 0.885 ± 0.291, as the model correctly identified absent or minimal necrotic cores as empty, though cases with small necrotic regions showed greater prediction variability.

**Figure 5.**
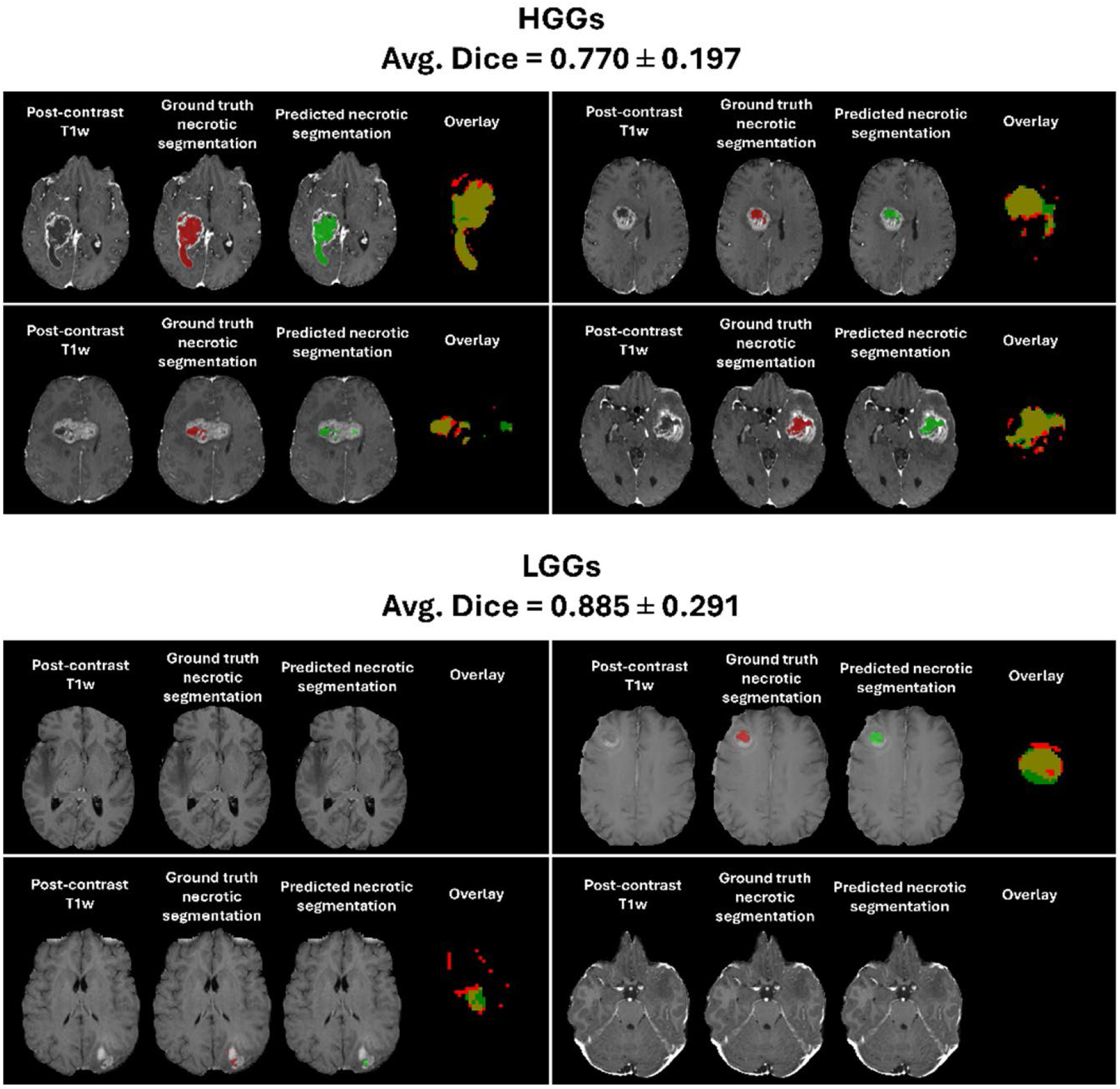
Qualitative evaluation of Stage 1 nnU-Net necrotic core segmentation performance for representative HGG and LGG cases. For each case, the post-contrast T1w image is shown alongside the ground truth necrotic segmentation (red), the predicted necrotic segmentation (green), and their overlay, where yellow indicates overlap between ground truth and prediction. Average Dice scores are reported for HGGs (0.770 ± 0.197) and LGGs (0.885 ± 0.291).

Quantitative results for the internal and external test sets, stratified by tumor type, are presented in Table 2. Across both internal tumor types and the external HGG dataset, whole-brain and whole-tumor metrics were equivalent between the model with and without necrotic scaling, though paired t-tests revealed small but significant differences in several brain-level metrics for internal HGG (SSIM, PSNR, and NMSE all significant). Internally, enhancing tumor Dice was higher with necrotic scaling than without, from 0.793 to 0.816 for HGGs (TOST p = 0.0649; t-test p = 0.3756) and from 0.219 to 0.451 for LGGs (TOST p = 0.8529; t-test p = 0.3264), though neither difference reached significance. The lower LGG values reflect that few internal LGG cases enhanced at all, and those that did had small enhancing regions, to which Dice is inherently more sensitive than HD95. Consistent with this, enhancing tumor HD95 showed no significant difference in the internal dataset for either tumor type, supporting that prediction quality was preserved despite the lower Dice scores in LGGs. On the external HGG dataset, necrotic scaling significantly improved enhancing tumor Dice from 0.465 to 0.612 (TOST p = 0.9900) and significantly reduced HD95 from 9.373 to 4.075 mm (TOST p = 0.9916), mirroring the Dice improvement in the same cohort.

### 3.3 Comparison with State-of-the-Art Models

Quantitative comparison with HRNet and DDResUNet is presented in Table 3. Across the internal HGG and LGG test sets, NeCS-Flow achieved the best performance for all evaluated metrics, with the largest gains for enhancing tumor segmentation. In LGGs, NeCS-Flow achieved an enhancing tumor Dice of 0.451, significantly exceeding HRNet (0.017) and DDResUNet, which predicted no enhancing-tumor voxels in any LGG case. In HGGs, NeCS-Flow achieved 0.816 versus 0.567 (HRNet) and 0.010 (DDResUNet). On the external HGG test set, all three models showed reduced performance relative to internal testing, consistent with domain shift. The external standard deviation (±0.07) was comparable to the internal one (±0.08), indicating a systematic reduction rather than failure in a subset. NeCS-Flow nonetheless outperformed both benchmarks on every metric, including enhancing tumor Dice (0.612 vs. 0.487 for HRNet and 0.000 for DDResUNet). HD95 further distinguished NeCS-Flow: internally, it achieved a significantly lower HD95 than HRNet (3.017 vs. 8.577 mm), while DDResUNet’s predictions were too sparse to yield a defined HD95. Externally, NeCS-Flow again achieved a significantly lower HD95 than HRNet (4.075 vs. 9.165 mm). Representative examples from the external GBM test set are provided in Supplementary Figure S2. Together, these findings show that NeCS-Flow’s advantage over existing methods is retained under domain shift.

### 3.4 Reader Assessment

The results of the readers’ discrimination between synthetic and real images are summarized in Table 4. Overall mean discrimination accuracy was 61.7%, modestly above chance. Performance varied across readers: one performed at chance (R1, 47.5%, p = 0.875), while the other two distinguished synthetic from real images at rates significantly above chance (R2, 70.0%, significant; R3, 67.5%, significant). Discrimination accuracy was non-significant for both tumor types (HGG and LGG: 61.7%, p = 0.092). Specificity exceeded sensitivity (66.7% vs. 56.7%), indicating that synthetic images were more frequently mistaken for real than the reverse. Among synthetic cases, higher enhancement realism ratings were associated with a greater likelihood of being misclassified as real (point biserial r = -0.420, p = 0.006).

**Table 4:** Real vs. synthetic discrimination analysis results.

|  | Result |
| --- | --- |
| *Per-Reader Discrimination Accuracy* |  |
| R1 | 47.5% (p=0.875) |
| R2 | 70.0% (p=0.017) |
| R3 | 67.5% (p=0.038) |
| Mean across readers | 61.7% |
| *Overall Sensitivity and Specificity* |  |
| Sensitivity (synthetics correctly identified) | 56.7% |
| Specificity (reals correctly identified) | 66.7% |
| *Discrimination Accuracy by Tumor Type* |  |
| HGG | 61.7% (p = 0.092) |
| LGG | 61.7% (p = 0.092) |
| *Enhancement Realism vs. Correct Identification* |  |
| Point-biserial correlation (synthetic cases only) | r = -0.420, p = 0.006 |

Per-tumor-type reader performance results are reported in Table 5A. No statistically significant differences between real and synthetic images were found for any metric, pooled or by tumor type. Diagnostic accuracy was equal (73.3% vs. 73.3%, p = 1.000), and image quality ratings were comparable (3.98 vs. 4.23, p = 0.177). Artifacts were detected less frequently in synthetic images (1.7% vs. 10.0%, p = 0.114). HGG cases showed closer real-synthetic parity across most metrics, while LGG cases showed the largest gap in enhancement detection accuracy (93.3% vs. 73.3%, p = 0.080). Reader consistency (Table 5B) was moderate for diagnosis (κ = 0.590) and substantial for enhancement detection (κ = 0.762). HGG cases showed stronger consistency across all metrics, while LGG cases showed lower consistency, with only 55.6% of enhancement realism ratings within ±1 point and a mean absolute difference of 1.22.

**Table 5A:** Reader study results stratified by tumor type.

| Metric | Tumor Type | Real | Synthetic | p-Value |
| --- | --- | --- | --- | --- |
| Diagnostic Accuracy (%) | Overall | 73.3% | 73.3% | 1.000 |
|  | HGG | 83.3% | 76.7% | 0.748 |
|  | LGG | 63.3% | 70.0% | 0.785 |
| Diagnostic Confidence (1-5) | Overall | 4.20 ± 1.10 | 3.95 ± 1.02 | 0.199 |
|  | HGG | 4.40 ± 0.72 | 4.10 ± 0.99 | 0.187 |
|  | LGG | 4.00 ± 1.36 | 3.80 ± 1.03 | 0.524 |
| Enhancement Detection (%) | Overall | 96.7% | 86.7% | 0.095 |
|  | HGG | 100.0% | 100.0% | 1.000 |
|  | LGG | 93.3% | 73.3% | 0.080 |
| Enhancement Realism (1-5) | Overall | 4.40 ± 0.89 | 4.10 ± 1.08 | 0.154 |
|  | HGG | 4.67 ± 0.61 | 4.40 ± 0.93 | 0.194 |
|  | LGG | 3.75 ± 1.14 | 3.33 ± 1.07 | 0.366 |
| Image Quality (1-5) | Overall | 3.98 ± 1.07 | 4.23 ± 0.95 | 0.177 |
|  | HGG | 4.30 ± 0.84 | 4.63 ± 0.49 | 0.065 |
|  | LGG | 3.67 ± 1.18 | 3.83 ± 1.12 | 0.577 |
| Artifacts Detected (%) | Overall | 10.0% | 1.7% | 0.114 |
|  | HGG | 3.3% | 0.0% | 1.000 |
|  | LGG | 16.7% | 3.3% | 0.195 |

Continuous metrics are reported as mean ± standard deviation.

**Table 5B:**
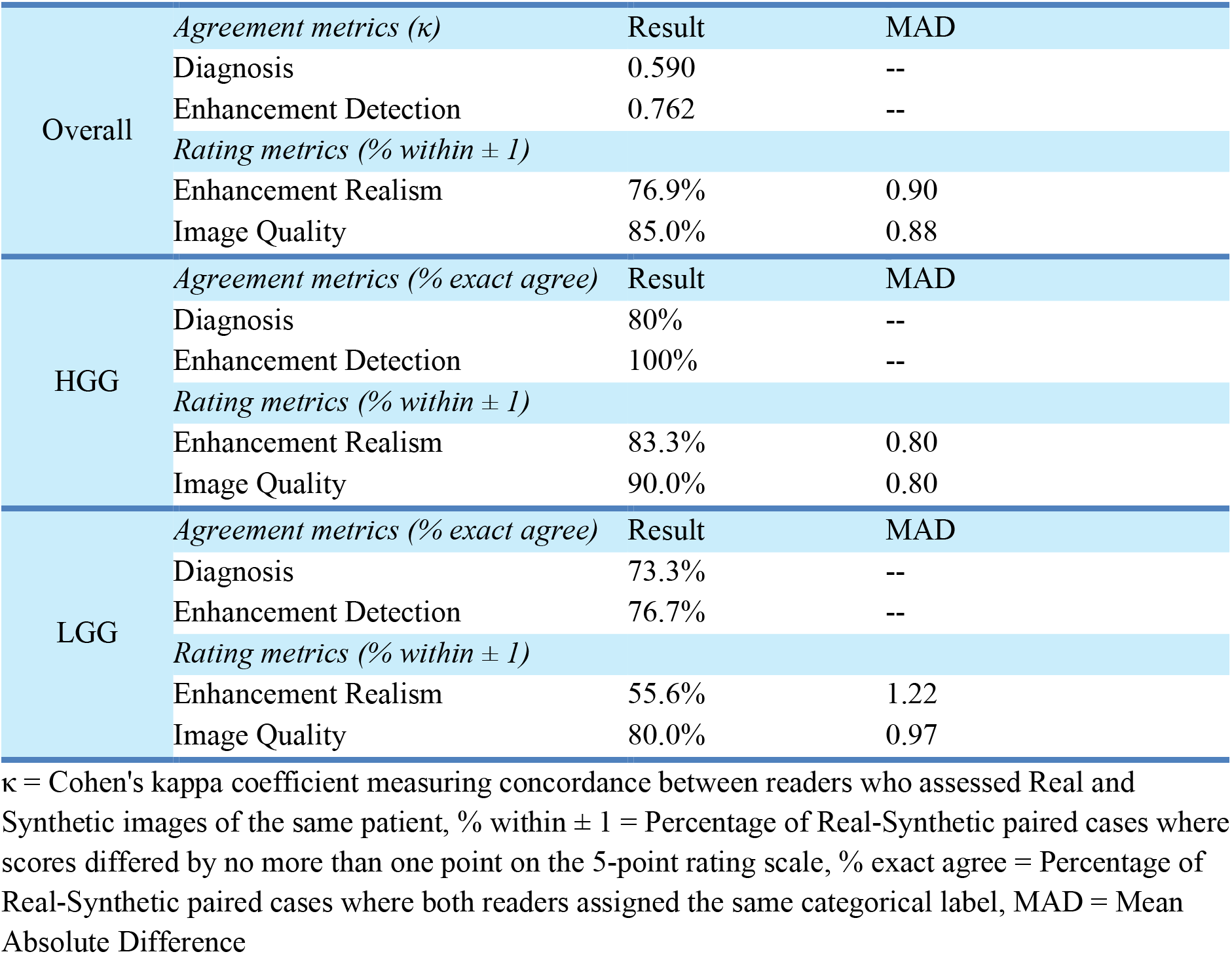
Reader consistency and real-versus-synthetic concordance analysis.

## 4. DISCUSSION

This study presents NeCS-Flow, a three-stage pipeline combining necrotic scaling and PMRF refinement for gadolinium-free post-contrast brain MRI synthesis. Necrotic scaling improved predicted contrast enhancement accuracy, most notably externally, while preserving whole-brain and whole-tumor image quality. PMRF improved perceptual realism without sacrificing voxelwise fidelity. NeCS-Flow also outperformed HRNet and DDResUNet across image quality and enhancing tumor metrics, internally and externally.

The necrotic scaling mechanism addresses the heterogeneous, spatially localized nature of tumor enhancement, a challenge prior tumor-region-weighted loss functions and semantic feature-extraction modules do not fully resolve, since they treat the tumor as relatively homogeneous rather than modeling the relationship between the necrotic core and its enhancing rim. NeCS-Flow’s superior performance relative to HRNet and DDResUNet suggests that biologically informed spatial priors can improve enhancement synthesis beyond architectural modifications alone. While the necrotic core is most conspicuous in HGGs, the mechanism proved broadly applicable to LGGs: LGG cases with a detectable core receive the same scaling, while cases without a detected core receive no scaling, which itself signals the absence of necrosis and discourages false-positive enhancement in regions where it is physiologically unlikely. Consistent with this, enhancing tumor Dice improvements were observed across both tumor cohorts. Whole-brain image quality metrics showed statistically significant but clinically negligible differences with necrotic scaling, within the predefined equivalence margins, indicating the localized enhancement gains came without a meaningful trade-off in overall fidelity. The qualitative improvements observed with necrotic scaling, however, were not fully captured by Dice alone, which quantifies overlap but is insensitive to boundary completeness and shape fidelity. Enhancing tumor HD95, which more directly penalizes boundary deviation, better captured necrotic scaling’s advantage at the necrotic/enhancing interface. This divergence suggests overlap- and boundary-based measures should be interpreted together. PMRF, meanwhile, addressed the well-known tendency of voxelwise-optimized networks toward over-smoothing, with the FID and KID reductions confirming its effectiveness.

The ablation analysis highlights complementary roles for the individual input sequences. The inferior T2w+FLAIR performance indicates that pre-contrast T1w provides structural information not fully recoverable from the other sequences, consistent with prior ablation studies identifying T1w as among the most informative inputs for post-contrast synthesis (7, 8, 32). The strong T1w+T2w performance for whole-brain metrics suggests T2w contributes substantially to global anatomical fidelity, while the relative advantage of T1w+FLAIR within tumor regions suggests FLAIR provides complementary information on tumor-associated changes, including peritumoral abnormalities, that may improve localized synthesis.

Prediction uncertainty maps from Monte Carlo dropout showed consistently higher voxelwise variance in enhancing tumor regions and cortical CSF spaces across both cohorts. Elevated tumor-region uncertainty is expected, since contrast enhancement has no direct pre-contrast correlate. Elevated CSF uncertainty is consistent with variable signal from CSF pulsation and flow effects. These estimates may support future confidence-aware synthesis frameworks, pending further work on clinical interpretation.

The reader study supports the framework’s clinical feasibility. The modest overall discrimination performance, and the finding that synthetic images were more frequently mistaken for real than the reverse, suggest that the PMRF stage plays a critical role in bridging the perceptual gap that has limited the clinical acceptance of synthesized images. This interpretation is reinforced by the significant negative correlation between enhancement realism and correct identification, indicating that the more realistic a synthetic case’s enhancement appeared to a reader, the more likely that reader was to misclassify it as real. This is consistent with PMRF’s perceptual refinement contributing to the visual realism of synthetic enhancement in a way that affects readers’ ability to distinguish synthetic from real images. The absence of significant differences in any clinical metric further suggests that the pipeline preserves diagnostically relevant enhancement information sufficiently to support radiological assessment.

Post-study interviews revealed meaningful differences in how readers approached image quality assessment. One reader evaluated image quality globally, focusing on cortical vessel visibility, gray-white matter contrast, and overall image sharpness, while the other two focused primarily on the tumor region. One reader noted that skull-stripping removed familiar anatomical context, complicating assessment of intraparenchymal and leptomeningeal enhancement for both real and synthetic images and occasionally lowering realism ratings unrelated to synthesis quality. Readers identified enhancing-rim completeness and peritumoral edema as primary indicators of realistic enhancement, particularly in HGG cases—features that should be specifically targeted in future synthesis work. The challenges in LGG cases reflect diagnostic ambiguity inherent to this tumor type: minimally enhancing tumors are almost invariably classified as LGG clinically, so several LGG cases with enhancement were misclassified as HGG, and subtle or focal enhancement is inherently subjective to assess, contributing to lower and more variable detection accuracy.

Performance on the external HGG cohort was lower than internally, but this reduction was not specific to the proposed method. HRNet showed a comparable decline, and DDResUNet, which produced negligible enhancement internally, produced no detectable enhancing tumor overlap externally. This is consistent with its original whole-brain loss without tumor-specific weighting, which underestimates small tumor regions relative to normal tissue (31). The relative ordering of the three methods was preserved, with NeCS-Flow’s Dice advantage over the best benchmark maintained externally, indicating the reduction reflects domain shift rather than a limitation specific to necrotic scaling or PMRF. In fact, the enhancing tumor Dice improvement from necrotic scaling was larger externally than internally, suggesting the biologically motivated spatial prior transfers across institutions even where absolute synthesis accuracy does not, with its contribution greatest when the network is least well matched to the data. Enhancing tumor HD95 increased only modestly from internal to external testing, a smaller relative degradation than for Dice, and the external HD95 distribution was narrower than the internal one despite comparable Dice standard deviations between cohorts. Possible explanations include a more homogeneous or smaller external cohort, which per-case error analysis in a larger external validation cohort could help distinguish.

Additional test cases including HGGs with absent or minimal necrosis, for which necrotic scaling provides limited spatial guidance, are presented in Supplementary Figure S1 to further characterize the scope of model limitations.

### 4.1 Limitations

This study has several limitations. The training set was imbalanced across tumor types; future work should use multi-institutional cohorts with balanced representation. All datasets used were publicly available research cohorts; prospective validation on clinical data from diverse institutions remains an important next step. External validation was limited to a single cohort of 20 HGG cases, with no external LGG data available and no reader assessment of external synthetic images. Prospective multi-institutional validation, and evaluation of domain adaptation or fine-tuning strategies, will be necessary before clinical deployment. Finally, the reader study involved only three readers evaluating a limited number of cases, so future studies should use larger reader panels and more cases for robust conclusions about diagnostic value.

### 4.2 Conclusion

This work demonstrates that gadolinium-free synthesis of post-contrast brain MRI is achievable with high perceptual and diagnostic fidelity across gliomas. NeCS-Flow, combining necrotic-scaling-guided enhancement prediction with PMRF-based perceptual refinement, outperformed existing synthesis architectures on all evaluated metrics internally and externally. The reader study results represent a meaningful step toward the clinical translation of contrast-free brain tumor imaging.

### 4.3 Data Availability

All imaging data used in this study are publicly available: UPENN-GBM, BraTS-TCGA-LGG, and UCSF-PDGM via The Cancer Imaging Archive (https://www.cancerimagingarchive.net), and the external validation cohort via OpenNeuro (https://doi.org/10.18112/openneuro.ds007045.v1.0.1). Code for the NeCS-Flow pipeline, including the necrotic-scaling, synthesis, and PMRF refinement stages, is available from the authors upon request.

## Notes

**Grant Support:** This work was supported by NIH grants R01 HD112923, R01 CA292091, R01 CA282516, and R01 NS109439.

### Competing Interest Statement

The authors have declared no competing interest.

## REFERENCES

1. Falk Delgado A, Van Westen D, Nilsson M, et al.: Diagnostic value of alternative techniques to gadolinium-based contrast agents in MR neuroimaging—a comprehensive overview. Insights Imaging 2019; 10:84.

2. Tan AC, Ashley DM, López GY, Malinzak M, Friedman HS, Khasraw M: Management of glioblastoma: State of the art and future directions. CA Cancer J Clin 2020; 70:299–312.

3. Gulani V, Calamante F, Shellock FG, Kanal E, Reeder SB: Gadolinium deposition in the brain: summary of evidence and recommendations. Lancet Neurol 2017; 16:564–570.

4. McDonald RJ, McDonald JS, Kallmes DF, et al.: Intracranial Gadolinium Deposition after Contrast-enhanced MR Imaging. Radiology 2015; 275:772–782.

5. Prince MR, Zhang H, Zou Z, Staron RB, Brill PW: Incidence of Immediate Gadolinium Contrast Media Reactions. Am J Roentgenol 2011; 196:W138–W143.

6. McKinnon C, Nandhabalan M, Murray SA, Plaha P: Glioblastoma: clinical presentation, diagnosis, and management. BMJ 2021; 374:n1560.

7. Calabrese E, Rudie JD, Rauschecker AM, Villanueva-Meyer JE, Cha S: Feasibility of Simulated Postcontrast MRI of Glioblastomas and Lower-Grade Gliomas by Using Three-dimensional Fully Convolutional Neural Networks. RSNA 2021; 3.

8. Chen C, Raymond C, Speier W, et al.: Synthesizing MR Image Contrast EnhancementUsing 3D High-resolution ConvNets. ResearchGate 2022.

9. Eidex Z, Safari M, Qiu RLJ, et al.: T1-contrast enhanced MRI generation from multi-parametric MRI for glioma patients with latent tumor conditioning. Med Phys 2025; 52:2064–2073.

10. Gong E, Pauly JM, Wintermark M, Zaharchuk G: Deep learning enables reduced gadolinium dose for contrast-enhanced brain MRI. J Magn Reson Imaging 2018; 48:330–340.

11. Jayachandran Preetha C, Meredig H, Brugnara G, et al.: Deep-learning-based synthesis of post-contrast T1-weighted MRI for tumour response assessment in neuro-oncology: a multicentre, retrospective cohort study. Lancet Digit Health 2021; 3:e784–e794.

12. Xie H, Lei Y, Wang T, et al.: Magnetic resonance imaging contrast enhancement synthesis using cascade networks with local supervision. Med Phys 2022; 49:3278–3287.

13. Crete F, Dolmiere T, Ladret P, Nicolas M: The blur effect: perception and estimation with a new no-reference perceptual blur metric. Edited by Rogowitz BE, Pappas TN, Daly SJ. San Jose, CA, USA; 2007:64920I.

14. Pianykh OS, Pospelova K, Kamboj NH: Modeling Human Perception of Image Quality. J Digit Imaging 2018; 31:768–775.

15. Moya-Sáez E, de Luis-García R, Alberola-López C: Toward deep learning replacement of gadolinium in neuro-oncology: A review of contrast-enhanced synthetic MRI. Frontiers 2023; 2.

16. Hambardzumyan D, Bergers G: Glioblastoma: Defining Tumor Niches. Trends Cancer 2015; 1:252–265.

17. Hanif F, Muzaffar K, Perveen kahkashan, Malhi S, Simjee S: Glioblastoma Multiforme: A Review of its Epidemiology and Pathogenesis through Clinical Presentation and Treatment. Asian Pac J Cancer Prev 2017; 18.

18. Eidex Z, Safari M, Ding J, et al.: An efficient 3D latent diffusion model for T1-contrast enhanced MRI generation. Biomed Phys Eng Express 2026; 12:015075.

19. Ohayon G, Michaeli T, Elad M: Posterior-Mean Rectified Flow: Towards Minimum MSE Photo-Realistic Image Restoration. 2024.

20. Bakas S, Sako C, Akbari H, et al.: Multi-parametric magnetic resonance imaging (mpMRI) scans for de novo Glioblastoma (GBM) patients from the University of Pennsylvania Health System (UPENN-GBM). 2021.

21. Calabrese E, Villanueva-Meyer J, Rudie J, et al.: The University of California San Francisco Preoperative Diffuse Glioma MRI (UCSF-PDGM). 2023.

22. Bakas S, Akbari H, Sotiras A, et al.: Segmentation Labels for the Pre-operative Scans of the TCGA-LGG collection. 2017.

23. Elena Filimonova, Augusto Leone, Francesco Carbone, et al.: Real-world Multi-Institutional Glioblastoma MRI Dataset with Standardized Preprocessing, Expert-Validated Segmentation, and MGMT Promoter Profiling. 2025.

24. Isensee F, Jaeger PF, Kohl SAA, Petersen J, Maier-Hein KH: nnU-Net: a self-configuring method for deep learning-based biomedical image segmentation. Nat Methods 2021; 18:203–211.

25. Ronneberger O, Fischer P, Brox T: U-Net: Convolutional Networks for Biomedical Image Segmentation. 2015.

26. Ibrahim MA, Hazhirkarzar B, Dublin AB: Gadolinium Magnetic Resonance Imaging. In StatPearls. Treasure Island (FL): StatPearls Publishing; 2026.

27. Schiffmann R, Van Der Knaap MS: Invited Article: An MRI-based approach to the diagnosis of white matter disorders. Neurology 2009; 72:750–759.

28. Durand-Muñoz C, Flores-Alvarez E, Moreno-Jimenez S, Roldan-Valadez E: Pre-operative apparent diffusion coefficient values and tumour region volumes as prognostic biomarkers in glioblastoma: correlation and progression-free survival analyses. Insights Imaging 2019; 10:36.

29. Baehring JM, Fulbright RK: Diffusion-weighted MRI in neuro-oncology. CNS Oncol 2012; 1:155–167.

30. Romano A, Palizzi S, Romano A, et al.: Diffusion Weighted Imaging in Neuro-Oncology: Diagnosis, Post-Treatment Changes, and Advanced Sequences—An Updated Review. Cancers 2023; 15:618.

31. Osman AFI, Tamam NM: Contrast-enhanced MRI synthesis using dense-dilated residual convolutions based 3D network toward elimination of gadolinium in neuro-oncology. Journal of Applied Clinical Medical Physics 2023.

32. Bône A, Ammari S, Menu Y, et al.: From Dose Reduction to Contrast Maximization: Can Deep Learning Amplify the Impact of Contrast Media on Brain Magnetic Resonance Image Quality? A Reader Study. Invest Radiol 2022; 57:527–535.

