## Supplemental File for "NeCS-Flow: Necrosis-Guided Contrast-Enhanced MRI Synthesis with Rectified-Flow Refinement"

### Supplementary Material

#### 1. QUANTITATIVE METRIC CALCULATION

##### 1.1 Whole-Brain and Whole-Tumor Similarity Metrics

All quantitative evaluations were implemented in MATLAB (R2024a). SSIM, PSNR, and NMSE were computed between synthetic and real post-contrast T1w images, restricted to voxels within the brain mask for whole-brain metrics and within the whole-tumor mask for whole-tumor metrics. PSNR was computed using the standard formula $10 \cdot \log_{10}(1/MSE)$ with a maximum signal value of 1.0 following min-max normalization, in which both synthetic and real images were scaled to the intensity range of the real image within the masked region. NMSE was computed as the mean squared error between normalized images divided by the mean squared signal of the normalized real image. SSIM was computed on the full normalized 3D volumes with zero-padding outside the mask, using MATLAB's built-in SSIM function.

##### 1.2 Whole-Tumor and Enhancing-Tumor Dice Scores

Dice scores for whole-tumor and enhancing-tumor regions were computed using a pooled lesion-level approach. Instance-labeled segmentation masks were used, in which each individual lesion receives a unique integer label. For each ground-truth lesion, the best-matching predicted lesion was identified using greedy IoU matching with a minimum IoU threshold of 0.1. A ground-truth lesion was considered detected if its best-matching predicted lesion exceeded this threshold. The pooled Dice score was then computed jointly over all matched voxels across all detected lesions within a patient, using the formula $2|A \cap B| / (|A| + |B|)$, where A and B represent the union of all matched predicted and ground-truth lesion voxels, respectively. Whole-tumor pooled Dice was computed using instance masks of all tumor subregions, and enhancing-tumor pooled Dice was computed using instance masks of the contrast-enhancing region only. All metrics were computed per patient and summarized as mean and standard deviation across patients, stratified by tumor type.

##### 1.3 Enhancing-Tumor Hausdorff Distance

The 95th-percentile Hausdorff distance (HD95) was computed between the surfaces of the ground-truth and predicted enhancing-tumor masks, in voxel units. Surface voxels were extracted by morphological erosion (3×3×3 structuring element) and subtracting the eroded mask from the original binary mask. A Euclidean distance transform was computed for each surface, and each surface's voxels were sampled at the corresponding distance transform of the opposing surface to obtain the set of nearest-neighbor distances in both directions. HD95 was defined as the 95th percentile of the pooled bidirectional distance distribution. Pooled HD95 was computed using the same lesion-matching procedure described in Section 1.2.

#### 2. NECROTIC SCALING PIPELINE

The necrotic scaling pipeline applies a spatially adaptive intensity scaling to the necrotic core regions in the pre-contrast T1w, T2w, and FLAIR input images prior to synthesis. The pipeline operates on physical signal intensities. For each patient, the predicted necrotic core segmentation is first processed with morphological closing using a spherical structuring element of radius 2 voxels to connect spatially fragmented components. Connected components are then identified using 26-connectivity, and only components exceeding a minimum size of 10 voxels are included in the smooth scaling procedure; smaller isolated voxels receive a default scaling of 1.0. For each qualifying component, a distance transform is computed from the component boundary inward. Scaling weights are assigned as a smooth gradient that increases from 1.0 at the boundary to a maximum of 2.0 at the core, using a linear ramp over 50% of the component's maximum inward distance. Components that are too thin to support a spatial gradient (maximum inward distance less than 2.0 voxels) receive a uniform scaling of 1.5. The resulting scaling map is applied multiplicatively to the physical intensities within the necrotic mask.

#### 3. MODEL TRAINING DETAILS

##### 3.1 Stage 1: Necrotic Segmentation (nnU-Net) Training

The Stage 1 nnU-Net model was trained using nnU-Net v2's self-configuring 3D full-resolution pipeline with all default settings. Network depth, patch size, and batch size were determined automatically from the dataset fingerprint. The model was optimized using stochastic gradient descent with Nesterov momentum ($\mu = 0.99$) and weight decay of $3\times10^{-5}$, with an initial learning rate of 0.01 decayed according to a polynomial schedule, lr·(1−epoch/1000)^0.9. Network parameters were initialized using He (Kaiming) normal initialization. The loss function was the sum of Dice loss and cross-entropy loss, applied at each deep supervision output. Training was performed for the nnU-Net default of 1,000 epochs (250 iterations per epoch), with no early stopping. The final-epoch model checkpoint was used for inference. Data augmentation was applied on-the-fly using nnU-Net's default pipeline, including random rotations (up to ±30° per axis), scaling, elastic deformation, gamma correction, brightness and contrast adjustment, additive Gaussian noise, and mirroring. Five-fold cross-validation was used as configured by default.

##### 3.2 Stage 2: Synthesis U-Net Training

The Stage 2 synthesis U-Net (3D residual bottleneck U-Net architecture; three encoder/decoder levels with 3, 4, and 5 residual blocks per level; 32 base filters; 3×3×3 convolutions; leaky ReLU activation with batch normalization) was trained using the Adam optimizer ($\beta_1 = 0.9$, $\beta_2 = 0.999$, $\varepsilon = 1\times10^{-7}$, Keras defaults) with a constant learning rate of $1\times10^{-3}$ and a dropout rate of 0.3. The loss function combined mean absolute error with a gradient/edge-preservation term, equally weighted ($\alpha = 0.5$), to encourage sharper predictions. Training used 100×100×100-voxel 3D patches, a per-GPU batch size of 2 (effective batch size 4 across two mirrored GPU replicas), and mixed-precision (float16) computation. Patches in which fewer than 20% of label voxels were nonzero were excluded from training. Data augmentation consisted of a single random rigid rotation about the axial acquisition plane, with rotation angle drawn uniformly from 0–30° in a randomly chosen direction, applied jointly to the input images and labels. No other augmentation was applied. Training was performed for a fixed 25 epochs with no early stopping or validation-based model selection. Checkpoints were saved after every epoch, and the final-epoch checkpoint was used for all reported results. Data were split 90/10 at the patient level into training and validation sets using a fixed random seed.

**3.3 Stage 3: PMRF Refinement Stage Training**

The PMRF refinement stage was trained using pre-computed Stage 2 U-Net predictions as the flow source. For each training case, the U-Net posterior-mean prediction was perturbed with Gaussian noise ($\sigma = 0.1$) to produce a noisy source image $z_1$. A random timestep t was sampled uniformly from [0, 1], and the interpolated image $z_t$ was computed as $z_t = t \cdot y + (1-t) \cdot z_1$, where y is the ground-truth post-contrast T1w image. The target vector field was defined as $v = y - z_1$, representing the direction from the noisy source to the ground truth. Both the U-Net prediction and the ground truth were Z-score normalized using non-zero voxel statistics prior to training. A dropout rate of 0.1 was used. Because pre-computed Stage 2 predictions were used as the flow source, no additional data augmentation was applied during flow training. The flow model was trained to predict v given $z_t$ and t using a flow-matching mean squared error loss. A 3D time-conditioned U-Net was used as the architecture, with t provided as an additional input channel. Training used the Adam optimizer with a learning rate of $1\times10^{-4}$, a batch size of 4, and

90/10 training/validation split. The model was trained for 50 epochs on two NVIDIA RTX 2080 GPUs, with a learning rate decay schedule applied across epochs. Model checkpoints were saved at each epoch and the final checkpoint was used for inference. No early stopping or validation-based model selection was used. At inference time, the trained flow model was applied to refine Stage 2 U-Net predictions by integrating the learned vector field from the noisy source to the target distribution using 20 integration steps.

#### 4. READER STUDY PROTOCOL

A blinded, multi-reader study was conducted to evaluate the clinical utility and perceptual realism of synthetic post-contrast T1w images. Three readers participated: two board-certified neuroradiologists with 3 and 2 years of attending experience (R1 and R2), and a neuroradiology fellow in the first year of fellowship (R3). Readers independently reviewed images using a web-based platform (PACSbin) and recorded their responses using standardized digital forms (REDCap). The study included 60 cases balanced across tumor type: 30 HGGs and 30 LGGs. For each case, the real and synthetic post-contrast T1w images were reviewed in parallel by two different readers following a pre-defined assignment, with one reader reviewing the real image and the other reviewing the synthetic image of the same patient. Each reader was assigned 40 cases in total, comprising 20 HGG and 20 LGG cases presented in random order. Cases were presented without any indication of whether the image was real or synthetic. Inter-reader communication regarding assessments was prohibited to ensure independent evaluation. Readers were asked to complete their assessments within a 14-day period.

##### 4.1 Assessment Criteria & Scoring

###### 4.1.1 Diagnostic Utility

| Question | Answer | | | | |
|---|---|---|---|---|---|
| Based on the image appearance, is this case most likely to represent a high-grade glioma, low-grade glioma, or metastatic tumor on histology? | HGG | | LGG | | METS |
| How confident are you in your diagnosis? | 1 (Not confident at all) | 2 (Unsure) | 3 (Neutral) | 4 (Somewhat confident) | 5 (Extremely confident) |

###### 4.1.2 Enhancement Quality

| Question | Answer | | | | |
|---|---|---|---|---|---|
| Is there any pathological enhancement present in this image? | Yes | | | No | |
| How realistic is the pathological enhancement pattern in this image? | 1 (Clearly artificial) | 2 (Somewhat artificial) | 3 (Moderately realistic but questionable) | 4 (Realistic) | 5 (Very realistic) |

###### 4.1.3 Image Quality

| Question | Answer | | | | |
|---|---|---|---|---|---|
| Are there any artifacts or unusual features that impact interpretation? | Yes | | | No | |
| How would you rate the overall quality of this image? | 1 (Very Poor) | 2 (Poor) | 3 (Adequate) | 4 (Good) | 5 (Excellent) |

###### 4.1.4 Synthetic vs. Real Discrimination

| Question | Answer | |
|---|---|---|
| Do you think this is a real or synthesized image? | Real | Synthesized |

##### 4.2 Statistical Analysis

##### 4.2.1 Diagnostic Accuracy

Diagnostic accuracy was calculated as the percentage of correct tumor type classifications against the ground truth diagnosis for both real and synthetic images, overall and stratified by tumor type. Differences between real and synthetic images were assessed using Fisher's exact test with a significance threshold of $p < 0.05$.

##### 4.2.2 Diagnostic Confidence, Enhancement Realism, and Image Quality

Mean scores on the 5-point rating scales for diagnostic confidence, enhancement realism, and overall image quality were compared between real and synthetic images using two-sample independent t-tests, overall and stratified by tumor type. Statistical significance was defined as $p < 0.05$. Enhancement realism ratings were restricted to cases where the reader indicated pathological enhancement was present.

##### 4.2.3 Enhancement Detection and Artifact Detection

Enhancement detection accuracy was calculated as the percentage of cases where the reader correctly identified the presence or absence of pathological enhancement, using ground truth enhancement status derived from tumor segmentation masks. Artifact detection rate was calculated as the percentage of cases where the reader reported interpretation-affecting artifacts. Both metrics were compared between real and synthetic images using Fisher's exact test with a significance threshold of $p < 0.05$.

##### 4.2.4 Real vs. Synthetic Discrimination

Discrimination accuracy was calculated as the percentage of cases correctly identified as real or synthetic per reader. P-values for per-reader and per-tumor-type discrimination accuracy were calculated using a binomial test against chance level (50%). Sensitivity was defined as the proportion of synthetic images correctly identified as synthetic, and specificity as the proportion of real images correctly identified as real. The point-biserial correlation assessed the association between enhancement realism scores and correct identification among synthetic cases only. A negative correlation indicates that higher realism scores were associated with lower likelihood of correct identification as synthetic.

##### 4.3 Inter-Reader Agreement Analysis

Given the parallel between-reader study design in which each patient was assessed by two different readers, one viewing the real image and the other viewing the synthetic image, concordance between real and synthetic image assessments for the same patients was evaluated as follows. For categorical variables (tumor type classification, enhancement presence detection), Cohen's kappa coefficient was computed as a measure of agreement between the reader who assessed the real image and the reader who assessed the synthetic image of the same patient. For continuous rating variables (diagnostic confidence, enhancement realism, image quality), the percentage of real-synthetic paired cases with scores differing by no more than one point on the 5-point scale was reported alongside the mean absolute difference between paired scores. All analyses were performed overall and stratified by tumor type.

#### 5. ADDITIONAL QUALITATIVE RESULTS

Figure S1 presents additional representative cases from the internal test set illustrating the effect of necrotic scaling on synthesis quality across both tumor types. Figure S2 presents representative cases from the external GBM test set, illustrating qualitative synthesis performance and generalization behavior relative to the internal test set.

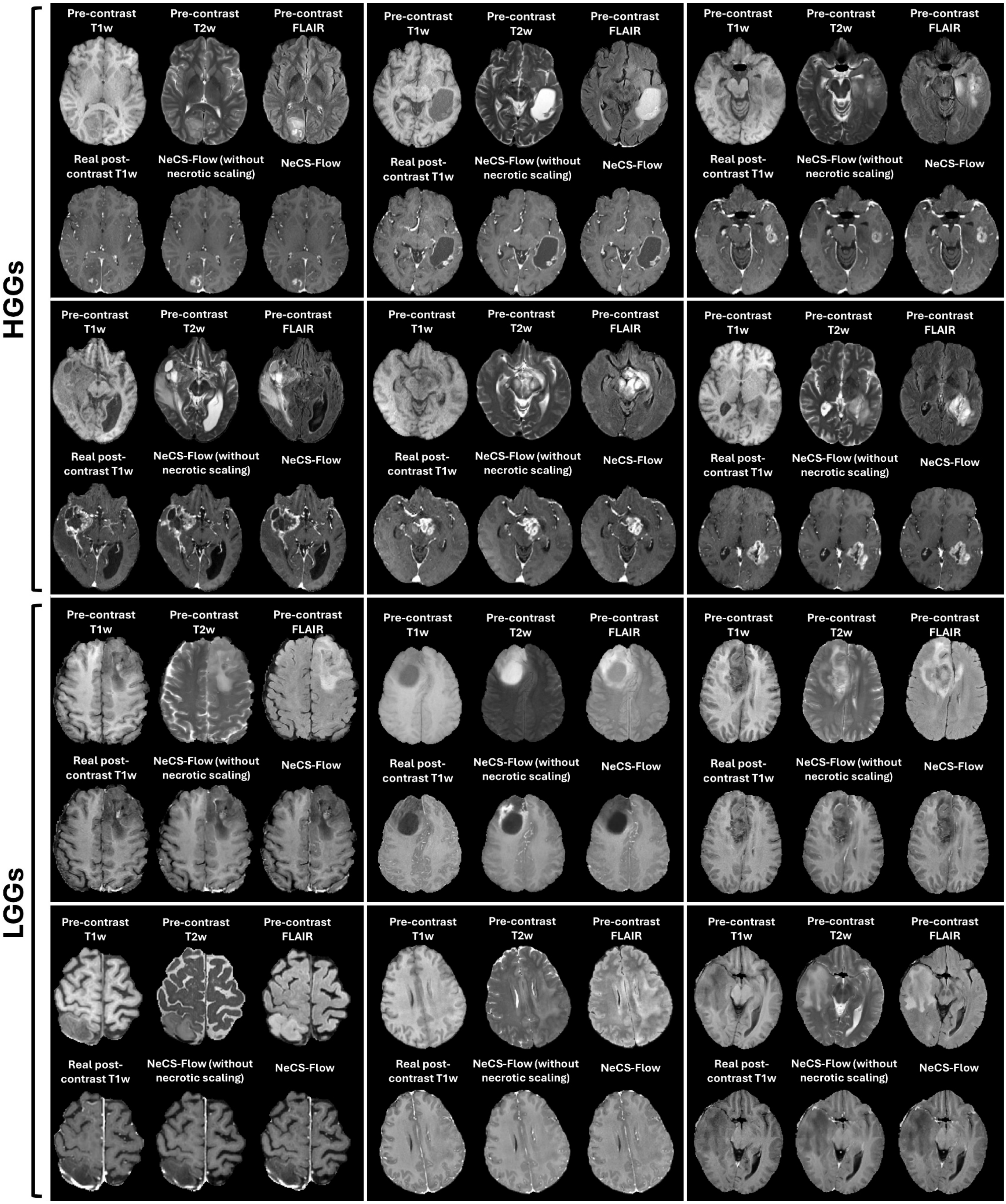
HGGs
LGGs
Pre-contrast T1w
Pre-contrast T2w
Pre-contrast FLAIR
Real post-contrast T1w
NeCS-Flow (without necrotic scaling)
NeCS-Flow

**Figure S1.** Qualitative comparison of predicted post-contrast T1w images with and without necrotic scaling for representative HGG and LGG cases. For each case, the pre-contrast T1w, T2w, and FLAIR inputs are shown in the top row alongside the real post-contrast T1w image, the predicted post-contrast T1w image without necrotic scaling, and the predicted post-contrast T1w image with necrotic scaling at the bottom row.

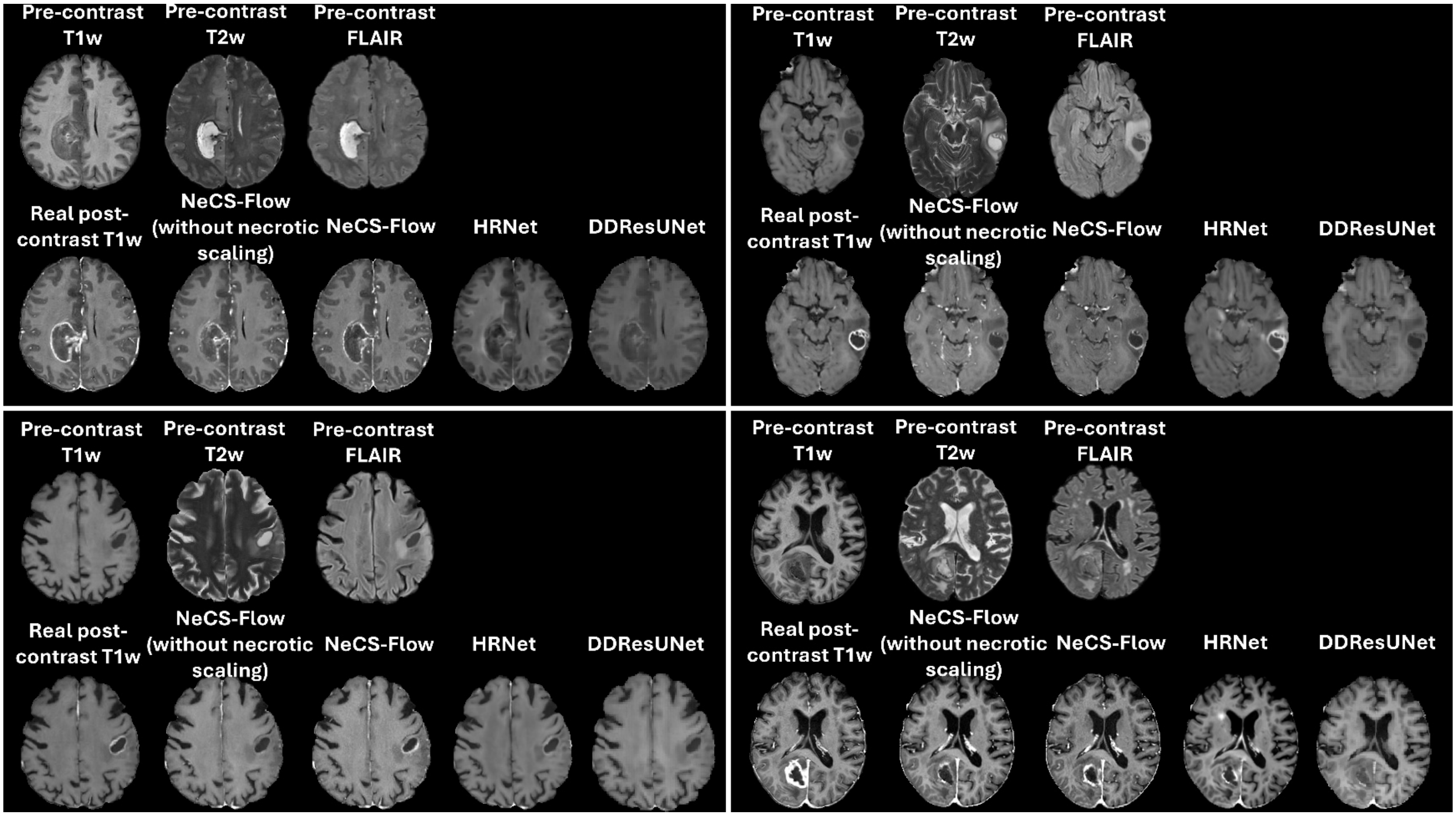


**Figure S2.** Qualitative comparison of predicted post-contrast T1w images across methods for four representative cases from the external GBM test set. For each case, the pre-contrast T1w, T2w, and FLAIR inputs are shown in the top row alongside the real post-contrast T1w image, the predicted post-contrast T1w image without necrotic scaling, the proposed method with necrotic scaling (NeCS-Flow), and two state-of-the-art baseline methods (HRNet and DDResUNet) in the bottom row.

#### 6. QUANTITATIVE RESULTS FOR ALTERNATIVE INTERNAL TEST SET

To assess the robustness of the quantitative results to the choice of test set, an alternative internal test set of 60 patients was constructed using a different random split of the same source datasets (30 HGG, 30 LGG). Supplementary Table S1 presents these quantitative results.

**Supplementary Table S1: Quantitative results of alternative internal test data stratified by tumor type.**

| Tumor Type | ROI | Metric | Without Necrotic Scaling | With Necrotic Scaling | TOST p-Value | t-test p-Value |
|---|---|---|---|---|---|---|
| HGG | Brain | SSIM | 0.964 ± 0.01 | 0.962 ± 0.01 | < 0.0001 | **< 0.0001** |
| | | PSNR (dB) | 28.24 ± 1.94 | 28.06 ± 1.89 | < 0.0001 | **0.0171** |
| | | NMSE | 0.041 ± 0.03 | 0.042 ± 0.03 | < 0.0001 | **0.0385** |
| | Whole Tumor | SSIM | 0.998 ± 0.00 | 0.998 ± 0.00 | < 0.0001 | **< 0.0001** |
| | | PSNR (dB) | 20.84 ± 2.46 | 20.69 ± 2.26 | 0.0001 | 0.7274 |
| | | NMSE | 0.107 ± 0.05 | 0.106 ± 0.03 | < 0.0001 | 0.9763 |
| | | Dice | 0.982 ± 0.01 | 0.983 ± 0.01 | < 0.0001 | 0.7417 |
| | Enhancing Tumor | Dice | 0.766 ± 0.15 | 0.796 ± 0.09 | 0.0016 | 0.6075 |
| | | HD95 (mm) | 2.268 ± 2.13 | 2.287 ± 1.60 | < 0.0001 | 0.9763 |
| LGG | Brain | SSIM | 0.963 ± 0.01 | 0.960 ± 0.01 | < 0.0001 | **< 0.0001** |
| | | PSNR (dB) | 26.66 ± 2.38 | 26.40 ± 2.29 | < 0.0001 | **0.0089** |
| | | NMSE | 0.034 ± 0.02 | 0.036 ± 0.02 | < 0.0001 | 0.0951 |
| | Whole Tumor | SSIM | 0.998 ± 0.00 | 0.998 ± 0.00 | < 0.0001 | **0.0429** |
| | | PSNR (dB) | 22.56 ± 3.36 | 22.20 ± 3.60 | 0.0214 | 0.5844 |
| | | NMSE | 0.050 ± 0.04 | 0.054 ± 0.04 | < 0.0001 | 0.6075 |
| | | Dice | 0.955 ± 0.10 | 0.971 ± 0.05 | < 0.0001 | 0.6075 |
| | Enhancing Tumor | Dice | 0.216 ± 0.00 | 0.422 ± 0.09 | **0.9012** | 0.0501 |
| | | HD95 (mm) | 10.33 ± 3.56 | 5.203 ± 1.63 | **0.6444** | 0.6075 |

SSIM = Structural Similarity Index Measure, PSNR = Peak Signal-to-Noise Ratio, NMSE = Normalized Mean Squared Error, HD95 = 95th-percentile Hausdorff distance, ROI = Region of Interest
